# Tracking changes in stability of North Sea Atlantic cod in 40 years

**DOI:** 10.1101/2025.01.06.631473

**Authors:** Hsiao-Hang Tao, Chih-hao Hsieh, Manuel Hidalgo, Vasilis Dakos

**Affiliations:** Institut des Sciences de l’Evolution de Montpellier, Université de Montpellier, Montpellier, France; Institute of Oceanography, National Taiwan University, Taipei 10617, Taiwan; Institute of Ecology and Evolutionary Biology, Department of Life Science, National Taiwan University, Taipei 10617, Taiwan; Research Center for Environmental Changes, Academia Sinica, Taipei 11529, Taiwan; National Center for Theoretical Sciences, Taipei 10617, Taiwan; Spanish Institute of Oceanography (IEO, CSIC), Balearic Oceanographic Center, Group of Ecosystem Oceanography (GRECO), Moll de Ponent s/n, 07190, Palma, Spain

**Keywords:** Age-specific abundance, non-linear dynamics, empirical dynamical modelling, interaction Jacobians, population stability, recovery dynamics

## Abstract

Tracking changes in the stability of low-abundance populations is vital for conservation. While the stability of natural populations is often assessed based on linear dynamics, many exhibit state-dependent dynamics, such as the iconic Atlantic cod in the North Sea (*Gadus morhua*). This fish population experienced an abrupt decline in the year 2000 and has not shown a clear sign of recovery despite relaxed fishing pressure. To understand the mechanisms behind the recovery dynamics, we used a dynamical systems approach to reconstruct the state space of the population from yearly abundance data of age groups between 1982-2021. Specifically, we estimated yearly interaction matrices of age groups and derived yearly population stability, defined as the rate of recovery to or departure from the current state after a small perturbation to the matrix. We found that since the abrupt decline in abundance of North Sea cod in the year 2000, the population continued to fluctuate closer to the transition threshold; some age groups experienced weaker self-regulation; and age-1 group became more sensitive to perturbations than older groups. These results provide important information on North Sea cod’s resilience and capacity to buffer natural and anthropogenic impacts.

## Introduction

To date, many marine populations are experiencing delayed rebuilding and uncertainties in recovery (Neubauer et al. 2013, Britten et al. 2017, Duarte et al. 2020). Understanding these populations’ stability, defined as the capacity of the populations to persist or maintain their present state in the face of disturbance, may provide explanations for their recovery dynamics. The stability of a fish population is often evaluated by the distance or rate of recovery to fixed biological reference points (e.g., Neubauer et al. 2013, Trochta et al. 2020). These reference points are determined by assessments of populations’ state based on linear dynamics. However, population dynamics are shaped by deterministic and stochastic forces (Bjørnstad and Grenfell 2001), and some marine fishes exhibit nonlinear dynamics with varying equilibrium points over time (Anderson et al. 2008, Glaser et al. 2014) and regime shift-like dynamics (Blöcker et al. 2023).

The mismatch between linear methods and nonlinear dynamics of natural systems motivates recent advances to estimate stability as how the system’s state departs from or recovers to the current state when slightly perturbed, without having to identify a reference or equilibrium point (Bjørnstad and Grenfell 2001, Ushio et al. 2018, Cenci and Saavedra 2019, Grziwotz et al. 2023). Particularly, empirical dynamic modeling, a statistical approach which is based on nonlinear dynamical systems (Takens 1981), has been used to estimate the stability of natural communities (Cenci and Saavedra 2019, Chang et al. 2021, Zhao et al. 2023), evaluate species interactions (Ushio et al. 2018), and rank species’ relative sensitivity to perturbations (Medeiros et al. 2022, Medeiros and Saavedra 2023). These properties are derived from reconstructed species interactions network within communities. However, whether empirical dynamic modeling can also estimate the stability of structured natural populations from interactions between age or size groups, remains unknown.

The interactions between age or size groups of a marine fish population fundamentally shape the dynamics and stability of the population (Hsieh et al. 2010, Hidalgo et al. 2011, Ciannelli et al. 2013, Wang et al. 2020). The most common interactions are growth, as positive influence from one age to the next; reproduction, as positive links from mature adult to young juveniles; cannibalism, as negative links from larger to smaller fish; and competition for food and habitats, as negative links from one age to another; self-regulation, as negative links within the same age groups. The interactions between age or size groups are strongly dependent on population abundance. At low abundance, adult reproduction is likely insufficient to sustain recruitment. If juvenile survival is further impacted by fishing pressure, growth within the life cycle is disrupted, resulting in fewer adults. In addition, all of the interactions change over time in the face of disturbances. Thus, the stability derived from age/size group interactions is not a stationary property yet changes with time.

Observing the change in the stability of a population through time provides potential explanations on how a natural population’s abundance or biomass changes in response to perturbation. In addition, the interactions between age/size groups and the sensitivity of age groups to perturbations provide additional information to better understand the recovery dynamics. This is particularly relevant to exploited fishes as fisheries are often age/size-selective (Rouyer et al. 2011, Hidalgo et al. 2011), and also to understand the recovery dynamics and mechanisms of depleted stocks.

Atlantic cod in the North Sea is an iconic example of fish experiencing delayed recovery. As a target commodity of one of the world’s oldest fisheries, North Sea cod have profoundly shaped the history, economy, and culture of Western Europe since the 16th century (Goodlad 2022). This population has experienced several abrupt changes in abundance and biomass for the past decades, with the most recent abrupt decline in 2000 (Blöcker et al. 2022), mainly due to substantial loss of juveniles (ICES 2024). Despite fishing pressure starting to decline in the same year, there was a lack of significant recovery. It remains unclear whether and how the stability of the population has changed, particularly before and after the events of abrupt decline in abundance and relaxed fishing pressure in 2000.

In this work, we used empirical dynamic modeling to study whether the failure in recover of Atlantic cod since 2000 is related to its stability and states of age groups. Using abundance time series of six age groups, we first recovered the trajectory of the population (manifold) in a reconstructed state space with six dimensions of age groups. We then estimated the pair-wise age group interactions at each time point based on locally-weighted linear regressions on the manifold (Sugihara et al. 2012). From these reconstructed interaction matrices, or the time-varying Jacobians which informs the local stability of the system, we extracted the dominant eigenvalues as a proxy of population stability. These eigenvalues describes the rate at which the system recovers to or diverges from its current state, after slightly perturbing the interaction networks (Ushio et al. 2018). We also extracted the dominant eigenvector of each age group from the interaction matrices to evaluate the relative sensitivity of an age group to perturbation in the age group interaction network at each time point (Medeiros et al. 2022).

We addressed the following questions: comparing two periods of 1982-2000 (before abrupt abundance decline) and 2001-2021 (after abrupt abundance decline), (1) how did the stability of the North Sea cod population differ? (2) what were the differences in the interactions among age groups? (3) what were the differences in the relative sensitivity of each age group to perturbations contributing to the population stability? We hypothesized that North Sea cod exhibited different stability patterns between the two periods. We further hypothesized several changes at the second period: due to the sharp decline in the population abundance, self-regulations within age groups were weakened. In addition, due to extreme low abundance of adults, reproduction success was reduced, resulting in higher sensitivity of younger groups to perturbations in the age group interaction network. In contrast, declines in the survival rate of younger groups impaired the growth in the life cycle, leading to less sensitivity of older groups to perturbations in the network.

## Methods

### 1. Data of time series of North Sea Atlantic cod abundance

We obtained the winter abundance time series data (from January to March) between 1983-2022 of Atlantic cod in the North Sea, from the International Bottom Trawl Survey (IBTS) of the International Council for the Exploration of the Sea (ICES) (https://data.ices.dk). The data is catch per unit effort (CPUE), measured as the number of fish trawled adjusted to one hour of hauling and per survey rectangle (1° longitude by 0.5° latitude) for each age group across the North Sea. To calculate the abundance for each age group in the entire North Sea, we summed the CPUE data from all rectangles. We used change point analysis (Pélissié et al. 2024), to detected the year when an abrupt shift in the total abundance occurred over the study period. Estimated yearly fishing mortality for age-2 to age-4 groups was compiled from the ICES stock assessment report (ICES 2020).

### 2. Analytical framework

Empirical dynamic modeling (EDM) is a nonlinear statistical approach based on state space reconstruction (Takens 1981). Unlike traditional models that rely on predefined equations, EDM reconstructs the underlying dynamical system directly from observed time series data. In this framework, the state of a dynamical system is depicted as a specific location within a multivariate coordinate space, referred to as “state space”, where the axes represent causally linked variables or lagged terms of a variable. The system’s state evolves over time following its intrinsic dynamic rules, thereby forming a trajectory. The ensemble of these trajectories constitutes a geometric structure known as an attractor manifold, which elucidates the relationships between variables over time (Sugihara et al. 2012).

In this study, we used the following methods of empirical dynamical modelling to address each of our research questions: (1) simplex projection (Sugihara and May 1990) to determine the optimal embedding dimension for each age group, (2) S-map (Sugihara 1994) to quantify the nonlinearity of the population, (3) MDR S-map (Chang et al. 2021) to identify interactions between age groups. We then (4) derived the absolute value of the dominant eigenvalue of the interaction matrix (Ushio et al. 2018) estimated from the MDR S-map as a proxy of population stability, and finally (5) used the eigenvector ranking method (Medeiros et al. 2022) to identify the relative sensitivity of each age group to perturbations.

#### Step 1: Determine the optimal embedding dimension for each age group

We first used simplex projection (Sugihara and May 1990) to determine the optimal embedding dimension for each age group. This step is crucial for reconstructing the state space of a dynamical system. The embedding dimension refers to the optimal number of state variables, or the optimal number of time lags of a state variable, needed to reconstruct the manifold in the state space.

For each age group, we divided the time series into a library set and a prediction set. Within the library set, we created state-space reconstructions for different embedding dimensions ranging from 2 to 10. For each state-space reconstruction, we constructed simplexes, which are geometric polygons formed from neighboring points around a given point. These simplexes were then used to predict the future states of the system. Specifically, for each point in the state space, the future state was predicted based on the weighted average of the future states of the points that form the simplex. The weights were inversely related to the distances from the points to the vertex of the simplex.

To evaluate the prediction skill for each embedding dimension, we compared the predicted states to the observed states in the prediction set using root mean square error (RMSE). The embedding dimension with the best prediction skill, indicated by the lowest RMSE, was identified as the optimal embedding dimension.

#### Step 2: Identify the nonlinearity of the population

We used the S-map method to evaluate the nonlinearity, or state dependency of the population (Sugihara 1994, Hsieh et al. 2005). For each age group, we first reconstructed the state space using the abundance time series data of different time lags, where the number of time series was determined by the optimal embedding dimension identified in the previous step. In the S-map approach, a locally weighted linear regression model is fitted for each point in the state space, where closer points have a greater influence on the fit.

The weighting of neighboring points is controlled by a tuning parameter, θ, which determines the degree of state dependency. When θ = 0, all neighbors are equally weighted, resulting in a linear model. When θ is greater than 0, closer neighbors are given exponentially more weight, enabling the model to capture more local and potentially nonlinear dynamics. The fitted local linear models are then used to make predictions for the system’s future states.

We ran the S-map with various values of θ, ranging from 0 to 8, and determined the optimal θ based on the best forecast skill, identified by the minimum mean absolute error (MAE). If the prediction skill improves as θ increases from 0, the target age group is deemed nonlinear. Conversely, if the best predictive skill occurs at θ = 0, the target age group’s dynamics is linear.

We followed the analytical procedure outlined in (Clark and Luis 2019) to assess the significance of nonlinearity for each age group. We first calculated ΔMAE, which is the difference between MAE at θ = 0 and the minimum MAE. Next, we generated 1,000 null models using phase-randomized surrogates, which preserves the basic statistical properties of the time series such as autocorrelation. We then compared the ΔMAE from the original time series with the null distribution of ΔMAE. The dynamics is considered significantly nonlinear if ΔMAE is higher than the 95^th^ percentile of the null distribution.

#### Step 3: Construct an interaction matrix of age groups for each year

We quantified the strength and the direction of interaction between each pair of age groups through time by adapting the Multiview Distance Regularized S-map method (MDR S-map) (Chang et al. 2021). This approach is developed to study species interactions within a community, by reconstructing multiple low-dimensional state spaces with the optimal embedding dimension to determine neighborhood relationships among data points. These distances are then ensembled to determine the weights used in the multivariate S-map. The interaction Jacobian elements in the Jacobian interaction matrix of each time point indicate pairwise interaction strengths and directions.

In this study, we used six age groups to reconstruct the low-dimensional state space, with maximum embedding dimension of six. The interaction strength of age group *i* at year *t* on age group *j* in the following year is approximated as the interaction Jacobian *(∂A*_*j(t+1)*_ */ ∂A*_*i(t)*_), where *A*_*i(t)*_ is the abundance of age *i* at year *t*. We compiled all the pairwise interaction Jacobians between the six age groups to generate a 6 × 6 discrete-time Jacobian matrix that is interpreted as the interaction network of our six age groups. For self-interactions, defined as the change of any age group influences on itself *(∂A*_*i(t+1)*_ */ ∂A*_*i(t)*_), we subtract the value of interaction Jacobian by 1 (Miki et al. 2024).

#### Step 4: Compute the population stability at each year

From the interaction matrix of age groups of each year, we derived the absolute value of the dominant eigenvalue (|DEV|) to indicate the population’s stability of that year (after Ushio et al. 2018), known as the local Lyaponov stability. This proxy implies that when given a small perturbation at a given time to the interaction matrix, how fast the population returns to or departs from the current state, without having to identify whether the state is at equilibrium.

When |DEV| equals to 1, the population is at a transition threshold between stable and unstable states. When |DEV| is below 1, the population is stable as it tends to recover to the current trajectory from disturbances. The lower the value, the more stable the population is at that moment. Conversely, when |DEV| is higher than 1, the population is unstable, as it tends to depart from the current trajectory after a small disturbance. The higher the value, the more unstable the population is at that moment.

#### Step 5: Quantify the relative sensitivity of age groups to perturbations

To determine the relative sensitivity of age groups to perturbations, we adopted the eigenvector ranking method (Medeiros et al. 2022), originally developed to rank the relative sensitivity of different species within a community. In this method, the relative sensitivity is indicated by the absolute dominant eigenvector of each species, derived from time-varying Jacobian interaction matrix reconstructed using S-map. The principle lies in when given small perturbations to the interaction network, the distribution of abundance of a species expands or contracts approximately along the direction of the dominant eigenvector at a rate specified by the dominant eigenvalue (Medeiros et al. 2022). That is, the absolute value of the eigenvector of a species determines how much perturbed abundance would differ from the unperturbed abundance at the next time point. For a sensitive species, the perturbed abundance would have a large difference between perturbed and unperturbed abundance.

We applied this method by constructing the interaction matrix from six age groups with MDR S-map. To compare the relative sensitivity between age groups for a given year, we standardized the dominant eigenvector of each age by dividing the square root of the sum of square of all six eigenvectors of that year. Then, we used the absolute values to infer the relative sensitivity of age groups to perturbations. The original method used ranking rather than the absolute value of eigenvector due to the uncertainty of interaction Jacobian approximation (Medeiros et al. 2022). Therefore, we produced another result using the ranking of the absolute dominant eigenvector, which was compared with the result using absolute value.

## Results

We first examined the changes in abundance of Atlantic cod in the North Sea over the study period from 1982 to 2021. Using change point analysis, we detected an abrupt decline in the total abundance in the year 2000, with a drop of more than 60% of the total abundance (**Fig 1a-b**). This decline was mainly attributed to the decrease in the abundance of age groups 1 and 2. Fishing mortality for age 2-4 groups increased from 1982 to 2000, and started to decline since (**Fig S1**). However, despite relaxed fishing pressure, North Sea cod did not show significant recovery in the following 20 years.

**Fig 1.**
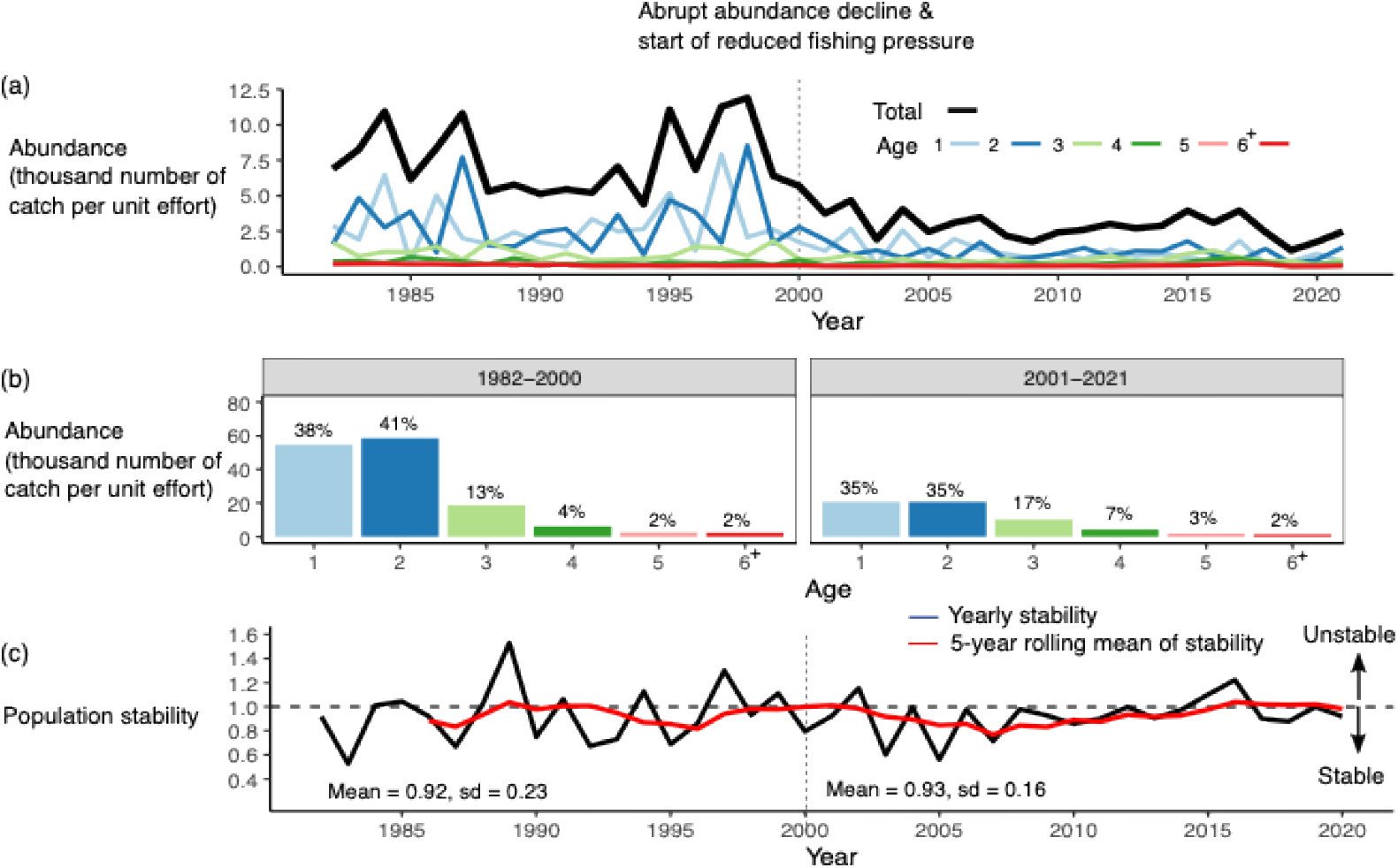
Changes in the abundance and stability of the North Sea cod population over time. (a) Black line indicates the abundance of the total population, and colored lines indicate abundances of age groups. Dotted vertical line indicates the year 2000, when the abundance of the population declined sharply, and fishing pressure started to decline in the same year. (b) Sum of yearly abundance for each age group within the period 1982-2000 and 2001-2021. The abundance of age groups 1 and 2 declined sharply from the first to the second period. The ratio above each bar indicates the percent abundance within each period (**c)** Black line indicates yearly population stability, computed as the absolute value of the dominant eigenvalue (DEV) from yearly Jacobian interaction matrix of age groups. The dotted black horizontal line indicates DEV = 1, when the population is at a transition period. A population is considered stable when DEV < 1 and unstable when DEV > 1. The red line indicates five-year rolling mean of stability. The mean and standard deviation of yearly stability values within each period are indicated. The mean DEV does not differ between 1982-2000 and 2001-2021; yet since year 2008, the population fluctuated closer to DEV = 1.

Next, using S-map, we found that North Sea cod exhibits nonlinear (disequilibrium) dynamics (**Fig S2**). This justified the need to use a nonlinear approach to estimate the stability of this population. To estimate time-varying stability of North Sea cod, we used MDR-Smap to estimate the age group interaction Jacobian of each year, and derived the absolute value of the dominant eigenvalue (|DEV|) as a population stability index. We found that throughout the study period, the population fluctuated between stable and unstable states (**Fig. 1c**). The mean |DEV| between the periods of 1983-2000 and 2001-2021 did not differ; yet during the second period, the fluctuations have become less pronounced (mean ± standard deviation for 1982-2000: 092±0.23; for 2001-2021: 093±0.16). Particularly, after year 2008, the population fluctuated close to the transition threshold (|DEV| = 1), imply that the population is approaching a transition.

We further compared the mean interaction strength of each age group pair between 1983-2000 and 2001-2021. The interaction strength of each age group pair and year were extracted from the corresponding Jacobian elements of the interaction matrix of that year. We averaged these yearly values to generate mean interaction strengths within each period. Overall, the strongest interaction in the network was the self-interaction link (how change in abundance of one age influences itself) (**Fig 2, S3**). The mean strengths of these links across periods were either very strong (|jacobian|>0.9) or strong (0.6 < |jacobian| <0.9). The second dominant interaction link in the network was the growth link (how change in one age influences the next age). The mean strengths of these links were either strong or medium (0.3 < |jacobian| <0.6). There was also a strong positive link from age-2 to age-1 group. All the other links were weak (|jacobian|<0.3).

**Fig 2.**
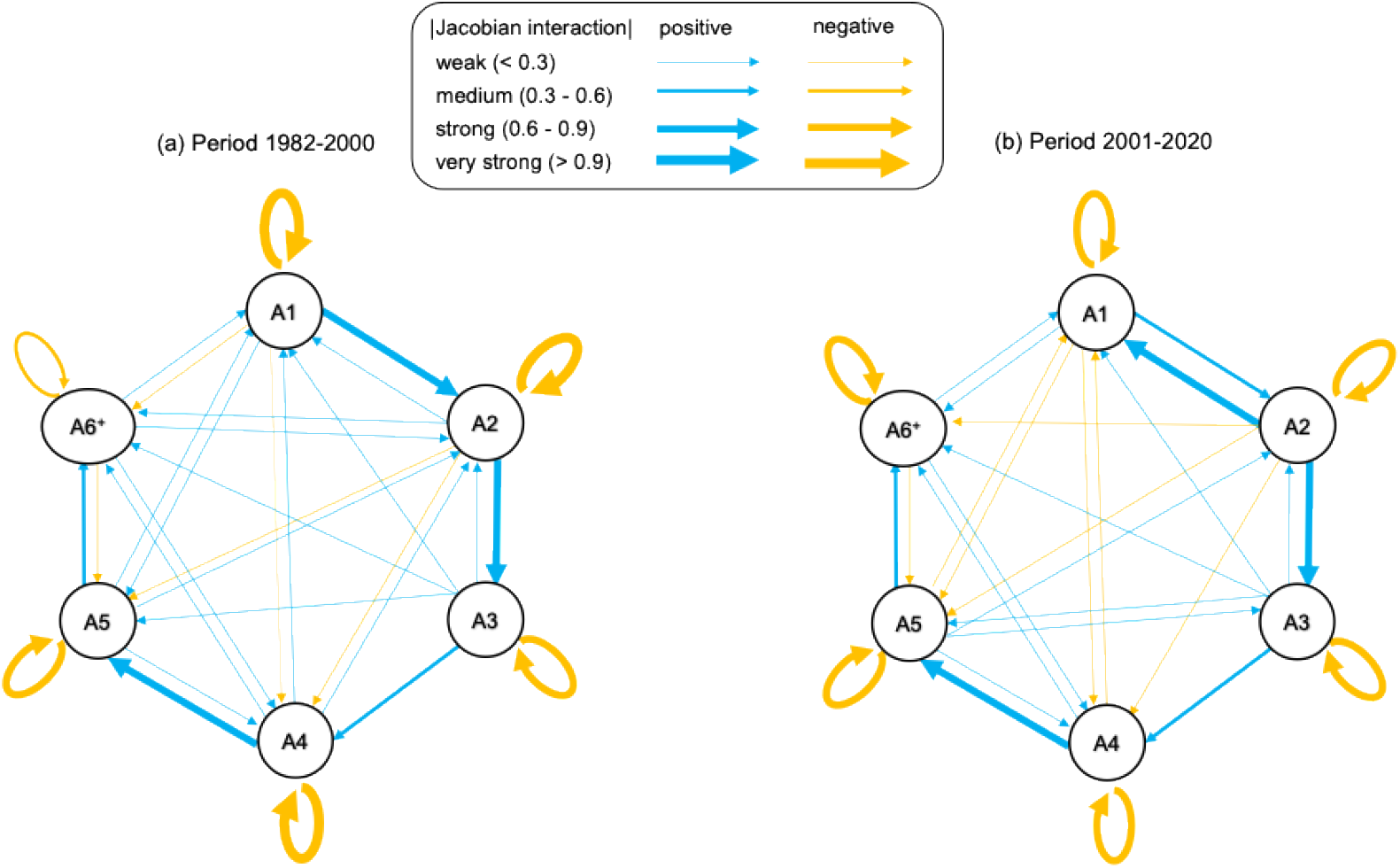
Interactions between age groups for the two time periods: 1982–2000 and 2001– 2021. The thickness of the arrows indicates the mean interaction strength, calculated as the average of yearly values for 1982–2000 (a) and 2021–2020 (b). Interaction strength is quantified as the absolute value of interaction Jacobian estimated from the MDR-Smap method. Here, we classified the interaction strength into four categories.

The major change in the age group interaction network between the two periods was that the self-interaction link within age-1, age-2, and age-4 groups decreased, while the link within age-6^+^ group increased (**Fig 2, S3**). In addition, the growth link from age-1 to age-2 group decreased. Conversely, the positive link from age-2 to age-1 increased.

Finally, we adopted the eigenvector ranking method to investigate the relative sensitivity of age groups. Comparing the distribution of these values between each age group within periods of 1983-2000 and 2001-2021 indicates that, before 2000, the relative sensitivity of different age groups was similar, although younger groups appeared to have slightly higher sensitivity (**Fig. 3**). In contrast, after 2000, the relative sensitivity of age-1 group to perturbations was much higher than other age groups, while the contribution of adult age groups (i.e. age ≥ 4 years) dropped. Using the rankings rather than the absolute values of the dominant eigenvector of the Jacobian interaction matrix showed comparable results (**Fig S4**). This implies that the contribution to stability was shared by the different age classes before 2000, while after 2000, the stability was mainly associated with the high sensitivity of the age-1 group.

**Fig 3.**
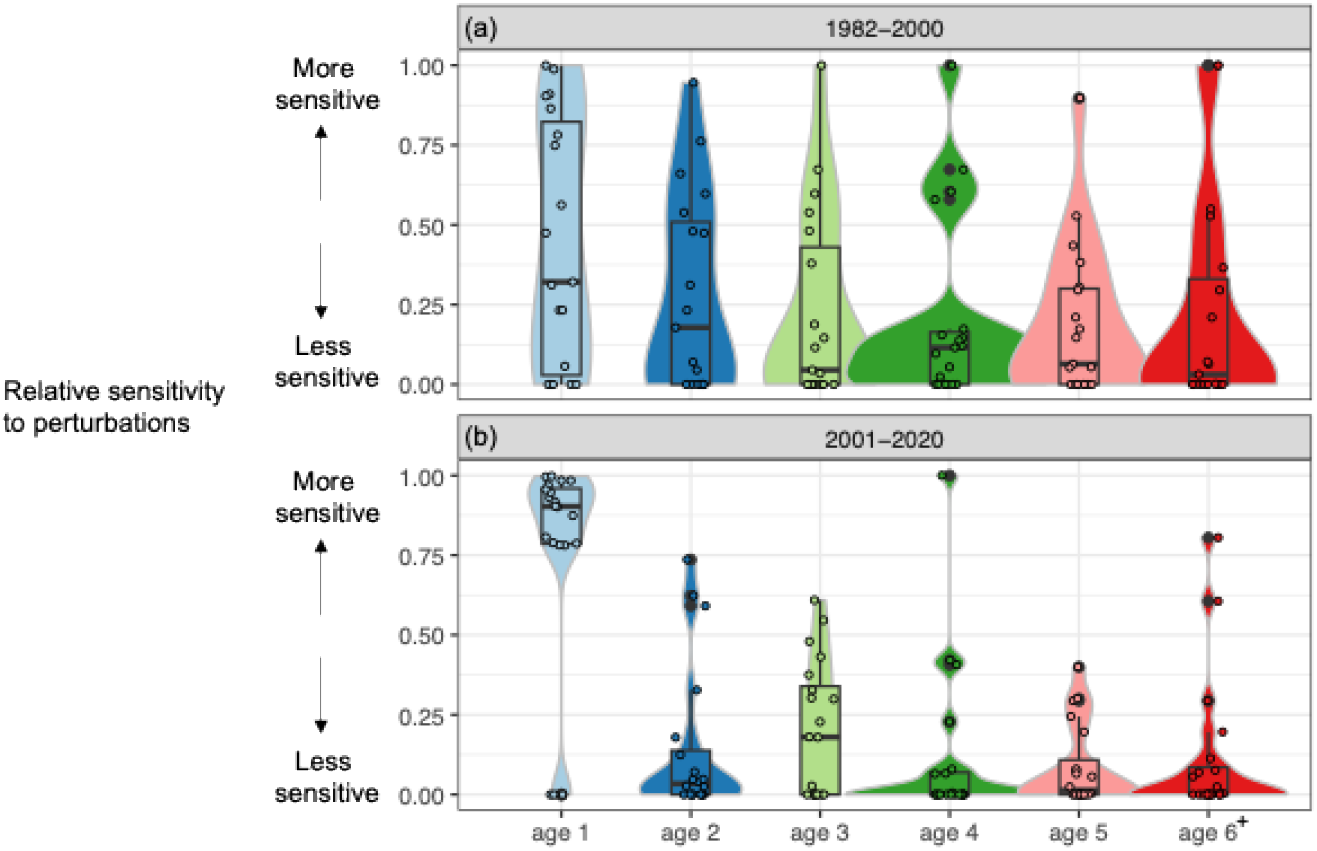
Distribution of the yearly relative sensitivity to perturbation of each age group for two periods 1983-2000 and 2001-2021. Each point represents the relative sensitivity of an age group compared to other age groups in a given year. The distribution of yearly relative sensitivity for each age group is summarized using boxplots and violin plots for the first period (a) and the second period (b). The relative sensitivity of each age group was computed as the absolute value of the leading eigenvector for each age group, standardized by the square root of the sum of squares of all six eigenvectors for a given year.

## Discussion

Despite reduced fishing pressure on many fisheries in recent decades, more than half of the assessed marine fish populations worldwide still need rebuilding (Worm et al. 2009). While fixed biological reference points are needed to estimate recovery and support management measures, exploited fish populations can exhibit nonlinear dynamics without fixed equilibrium (Glaser et al. 2014). This highlights the importance of accounting for state-dependent dynamics when evaluating the stability of these populations. In this work, we applied a dynamical systems method to investigate North Sea cod, an iconic fish population that suffered from an abrupt decline in 2000 and showed minimal recovery since, despite reduced fishing pressure. We analyzed and compared two time periods, 1982–2000 and 2001–2021, to evaluate (1) population stability, (2) interactions among age groups, and (3) the relative sensitivity of each age group to perturbations.

We found that the stability pattern of the North Sea cod appears to be different between the two periods. During the first period, the population strongly fluctuated between stable and unstable states (crossing frequently the transition threshold of |DEV| = 1, **Fig. 1c**). However, during the second period, particularly after 2008, the population mainly stayed at the stable zone (|DEV| < 1) and very close to the threshold (**Fig 1c**). Although it is hard to conclude if the cod population’s stability increased or decreased between the two periods, it seems that the change of the stability pattern followed the abrupt decline in abundance in year 2000.

One might argue that the low abundance during the second period could be the reason for the North Sea cod’s stability pattern, which stayed at the stable zone but close to the transition threshold. However, theoretical models show that even low-equilibrium states could be associated with high stability far away from the transition threshold (Dakos et al. 2017). Instead, such pattern of stability remaining close to yet below the transition implies several possibilities. In the best scenario, the population is going to cross the threshold and shift to a new state with higher abundance, as a sign of recovery. Alternatively, the population may shift to an attractor associated with lower abundance, implying further dampening of the population dynamics. Still, the population may move away from the threshold to a higher level of stability, meaning that the population will be “locked in” to its current low abundance state. Our method does not allow us to predict the direction and probability of change, yet the sign of North Sea cod approaching a transition threshold since year 2008 implies a change may occur, requiring attentions from fisheries management. Our finding can also complement other work focusing on forecasting, for example early warning signals (Clements et al. 2019, Grziwotz et al. 2023).

We found some clear differences in the age group interactions between the two periods, which may explain the recovery dynamics of this population. Particularly, after year 2000, the self-interaction strengths within both age-1 and age-2 groups decreased, while positive links from age-2 to age-1 group increased (**Fig. 2**). We speculated that these results were related to a positive density-dependent effect, where the sharp decline in juvenile’s abundance potentially reduces cannibalism and competition, leading to relaxed self-regulation to promoted survival of the juveniles (Hilborn et al. 2014, Perälä et al. 2022). In addition, the self-interaction link within age-4 group also decreased during the second period. This has important implications to population dynamics, because these older spawners can potentially contribute to larger and higher-quality eggs, promoting reproduction success. On the other hand, the growth interaction, defined as the positive links from one age to the next, were as strong as self-interaction links (**Fig. 2**). However, the strength of most growth interactions did not differ between the two periods. This suggests that even though relaxed self-regulation of age-1, age-2, and age-4 groups may have promoted their survival and abundance, these effects did not propagate to increase the strengh of growth toward the next age groups. This may explain why the North Sea cod has been trapped in the low-abundance state since year 2000.

In addition to age group interactions, we identified a major change in the relative sensitivity to perturbations among age groups between the two periods: age-1 group became much more sensitive to perturbations than older groups at the second period (**Fig. 3**). It is important to note that the sensitivity of age groups in this study is quantified as how sensitive the abundance of any age group depends on small perturbations in abundance of other age groups in the interaction network. In the context of a life cycle within a fish population, the abundance of juveniles mainly depends on the reproduction success of adults (with minor impact of cannibalism), while the abundance of adults relies the growth of younger groups. Based on these assumptions, the increased sensitivity of the age-1 group after year 2000 was potentially due to impaired reproduction from adult cod with their extremely low abundance. Another possible explanation is that age-1 group may have become more vulnerability to environmental forcings due to eroded demography, as documented in some exploited fishes (Anderson et al. 2008, Hsieh et al. 2010, Hidalgo et al. 2012). On the other hand, while age-1group became more sensitive to perturbations during the second period, the older groups became much less sensitive to perturbations (**Fig 3**). If our assumptions remain true, this result implies that the capacity to grow from one age to the next were impaired in the life cycle of this population, beginning from the sharp decline of age-1 group since year 2000. Indeed, we found that the strength of growth from age-1 to age-2 group was reduced since year 2000 (**Fig 2**), while the growth from age-2 onward did not change; yet the low abundance of age-1 group may have had a cascading effect on the entire growth chain. In summary, we speculated that during the second period, failures in reproduction from mature cod may have led to high sensitivity of age-1 group, while impaired growth chain may have led to reduced sensitivity of older age groups; these changes led to the population to be trapped in a low-abundance state until present.

This study has several limitations. First, we assessed the stability of the entire North Sea cod population, although recent evidence suggests it may comprise multiple subpopulations with potential mixing (Holmes et al. 2014). Additionally, our analysis was based on yearly winter data, as data from other seasons are shorter and less continuous. Winter is the season when mature cod undertake spawning migrations and aggregate spatially more than in other seasons (Neat et al. 2014). This spatial restructuring likely influences interactions among adults and between adults and juveniles. Another key limitation involves the data requirements for empirical dynamic modeling. At least 35 data points are typically needed to reliably reconstruct the state space (Sugihara et al. 2012), and the accuracy improves with longer, less noisy time series (Chang et al. 2017). This issue of the length of time series also affects other state-dependent approaches for assessing population stability, such as locally linear autoregressive state-space models for nonlinear dynamics (Ives and Dakos 2012), lagged time series reconstructions (Grziwotz et al. 2023), or stability proxies like the trace derived from interaction matrices (Cenci and Saavedra 2019). In this work, our time series was relatively short with 40 time points. We are therefore aware of potential process and observation errors when estimating |DEV| (Chang et al. 2021).

Future research can compare the available approaches to provide a comprehensive understanding of the stability of natural populations. Beyond the standard approach of estimating maximum sustainable yield in fisheries sciences (Lotze et al. 2011) and aforementioned state-dependent methods, several other techniques exist for quantifying the stability of marine fish populations. These include detecting abrupt shifts (Pélissié et al. 2024), identifying regime shifts (Blöcker et al. 2022), analyzing variations in time series (Hsieh et al. 2006), and detecting early warning signals (Clements et al. 2019). Each method defines stability differently with unique assumptions. A further systematic comparison of these methods could help clarify their applicability and improve the understanding of natural population dynamics across taxa and regions.

Finally, our approach of measuring population stability based on nonlinear dynamics, along with aforementioned methods in detecting regime shifts and early warning signals (e.g., Clements et al. 2019, Möllmann et al. 2021, Pélissié et al. 2024), may be considered to complement fisheries management advice with biological reference points obtained from state-of-the-art fisheries stock assessment methods and management strategy evaluations. These methods provide important information of the state of the populations beyond biomass-based sustainable indicators, and are more related with their resilience and capacities to buffer natural and anthropogenic impacts.

## Conclusion

Tracking changes in the stability of low-abundance populations is critical for understanding the mechanisms behind recovery dynamics. Using an analytical framework based on dynamical systems, we find that since the abrupt decline in abundance of North Sea cod in the year 2000, the population fluctuated closer to the transition threshold; some age groups experienced weaker self-regulation; and age-1 group became more sensitive to perturbations than older groups. We speculate that despite positive density-dependent effects to promote the survival of some age groups, impaired reproductive success from adults and reduced capacity to grow from one age to another resulted in a low-abundance state of cod since year 2000 without a strong sign of recovery. Our findings also demonstrate that the population appears to approach a transition since year 2008, yet is it unknown whether the abundance of the population is going to increase, decrease, or whether the age structure of the population will change. In any case, more attention in managing and preserving this population is important. Currently, commercial catches of the North Sea cod are predominantly composed of juveniles under three years old, with bycatch and discard rates estimated at 23% by weight of the total catch in 2023 (ICES 2024). Given the high sensitivity of juveniles to perturbations and their strong dependence on the recruitment of adults, it is critical to protect the entire population from overfishing and to preserve their habitats to ensure the sustainability of the population. The recovery of North Sea cod is also facing challenges in the rising ocean temperatures, increasing frequency and intensity of extreme events (IPCC 2023), and the potential irreversibility of regime shifts in the North Sea (Sguotti et al. 2022). Beyond Atlantic cod in the North Sea, several other exploited fish populations in this region also exhibited regime-shift dynamics (Blöcker et al. 2023). Our analytical framework is broadly applicable to other natural populations across taxa and regions with biomass or abundance time series for size, age, or life stages within the populations. More importantly, it enables the estimation of within-population interactions and sensitivities to perturbations, providing valuable insights to understanding recovery dynamics and informing conservation efforts.

## Code and data availability

All the data used this study were downloaded from open-access online databases as mentioned above. We used “rEDM” package (https://cran.r-project.org/src/contrib/Archive/rEDM, version 1.2.3) to perform all the analysis. The MDR S-map was performed using functions provided in Chang et al. (2021). All statistical analyses were performed in R 4.3.1. Data and codes for reproducing the results are deposited in Dryad https://datadryad.org/stash/share/J51GkKEfPNjlyiVhSccjYz-qDbtnx3O_1wbaK7CjXL4.

## Acknowledgements

We thank the research and survey teams of International Bottom Trawl Survey Working Group from International Council for the Exploration of the Sea (ICES) for providing the data. We thank Alejandro Cano, Benoît Pichon, and Ismaël Lajaaiti for commenting on an earlier version. This work is funded by Marie Curie postdoctoral fellowship of H.H-T. (HORIZON-MSCA-2021-PF-01, European Commission, LSP 244267; MAFIS project). C.H.H. is supported by the National Science and Technology Council, Taiwan.

