## supplementary materials for "Tracking changes in stability of North Sea Atlantic cod in 40 years"

**Title**

**Supplementary materials**

Fig S1-S4

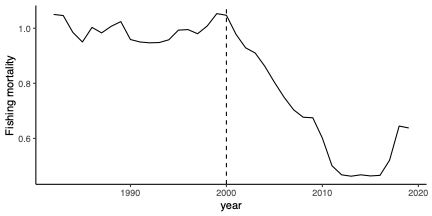

**Fig S1 Fishing mortality of North Sea cod**. The fishing mortality is estimated for age-2 to age-4 groups.

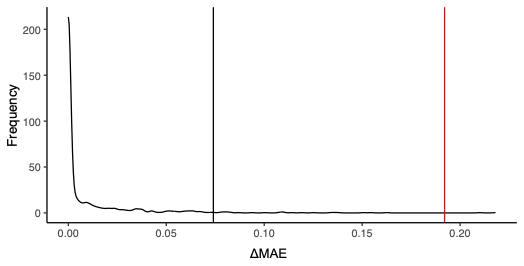

**Fig S2 Nonlinearity of the North Sea cod.** The nonlinearity of cod was determined by comparing the value of ΔMAE (0.19, red line) with the distribution of ΔMAE from null models (black curve) using S-map. The black vertical line indicates the 97.5% quantile of the null distribution. The population dynamics is considered nonlinear if ΔMAE is higher than the 97.5% percentile of the null distribution. We used phase-randomized surrogates to generate 1,000 null models.

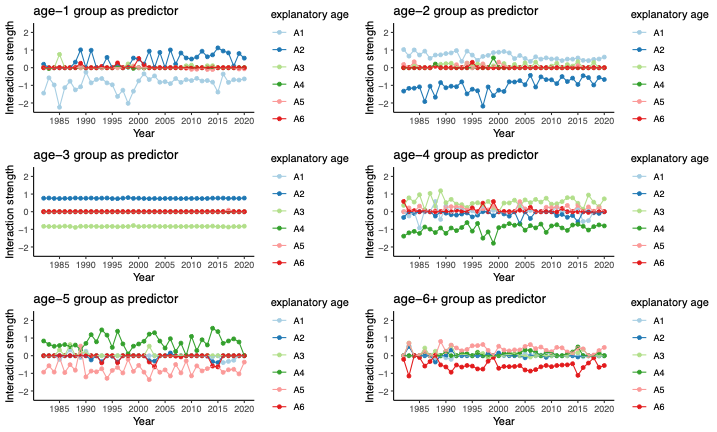

**Fig S3 Yearly interaction strengths between pairs of age groups during 1982-2021.** Interaction strength is the value of the interaction jacobian derived from the MDR S-map. Note that the interactions are directional: each explanatory age group (color) contributes to each predictor age group (panel). A1: age-1 group, A2: age-2 group, A3: age-3 group, A4: age-4 group, A5: age-5 group, A6: age-6^+^ group.

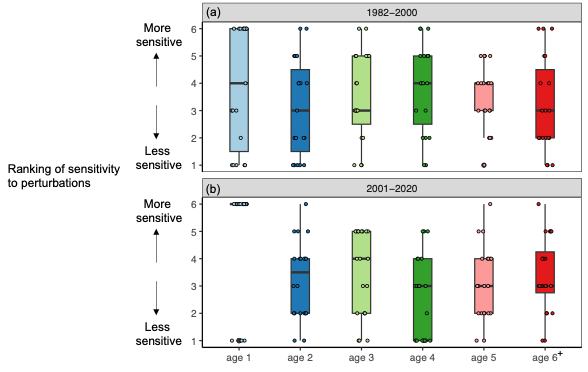

**Fig S4 Distribution of yearly ranking of sensitivity to perturbations of each age group for 1983-2000 and 2001-2021.** Each point represents the ranking of an age group compared to other age groups in a given year. The ranking scales from 1 (the least sensitive) to 6 (the most sensitive). The distribution of yearly ranking for each age group is summarized using boxplots plots for the first period (a) and the second period (b). Ranking of sensitivity to perturbations between age groups for a given year is computed as the ranking of the absolute value of dominant eigenvector between age groups from the jacobian interaction matrix, standardized by the square root of the sum of squares of all six eigenvectors for a given year, following the eigenvector ranking method in Medeiros et al. (2022).
